# A Continuous Attractor Model with Realistic Neural and Synaptic Properties Quantitatively Reproduces Grid Cell Physiology

**DOI:** 10.1101/2024.04.29.591748

**Authors:** Nate Sutton, Blanca Gutiérrez-Guzmán, Holger Dannenberg, Giorgio A. Ascoli

## Abstract

Computational simulations with data-driven physiological detail can foster a deeper understanding of the neural mechanisms involved in cognition. Here, we utilize the wealth of cellular properties from Hippocampome.org to study neural mechanisms of spatial coding with a spiking continuous attractor network model of medial entorhinal cortex circuit activity. The primary goal was to investigate if adding such realistic constraints could produce firing patterns similar to those measured in real neurons. Biological characteristics included in the work are excitability, connectivity, and synaptic signaling of neuron types defined primarily by their axonal and dendritic morphologies. We investigate the spiking dynamics in specific neuron types and the synaptic activities between groups of neurons. Modeling the rodent hippocampal formation keeps the simulations to a computationally reasonable scale while also anchoring the parameters and results to experimental measurements. Our model generates grid cell activity that well matches the spacing, size, and firing rates of grid fields recorded in live behaving animals from both published datasets and new experiments performed for this study. Our simulations also recreate different scales of those properties, e.g., small and large, as found along the dorsoventral axis of the medial entorhinal cortex. Computational exploration of neuronal and synaptic model parameters reveals that a broad range of neural properties produce grid fields in the simulation. These results demonstrate that the continuous attractor network model of grid cells is compatible with a spiking neural network implementation sourcing data-driven biophysical and anatomical parameters from Hippocampome.org. The software is released as open source to enable broad community reuse and encourage novel applications.

## Introduction

The hippocampus and surrounding circuits, including the entorhinal cortex, play a crucial role in representing and associating space, context, and episodic memory. Despite remarkable advances in characterizing the molecular and cellular properties of hippocampal neurons, a coherent framework to link this wealth of data with network-level computational function is still missing. Simulating cells involved in spatial cognition with detailed levels of physiological realism may foster mechanistic investigations of their roles in spatial navigation. Grid cells are place-modulated neurons with firing fields at multiple discrete and periodically spaced locations (Moser et al., 2014). These fields form a hexagonal pattern that tiles the entire space available to the animal. Many computational simulations have helped advance theories on how grid cells and other cellular activities are involved in spatial representation (Burak and Fiete, 2009; Hasselmo et al., 2007; Solanka et al., 2015; Zilli, 2012). Continuous attractor networks (CAN) are a major class of computational models used to recreate grid cell activities (Shipston-Sharman et al., 2016).

This study investigates whether the CAN model of grid cells is compatible with measured biophysical details of the underling neural network, such as neuron type-specific excitability, connectivity, and synaptic signaling. Finding an answer is important because it could add helpful realism to spatial navigation modeling. Prior work successfully modeled selected physiological characteristics (e.g., Pastoll et al., 2013), but obstacles remain in systematically finding relevant experimental measurements, designing a computationally efficient system that is compatible with those details yet can run at a useful neural scale, and acquiring animal data to compare modeled results. This work tackles those obstacles by taking advantage of existing open access resources to test a unique combination of biological details in a data-driven spiking neural network (SNN) model of pertinent portions of the entorhinal circuit. The results contribute supporting evidence in favor of the plausibility of the CAN model.

One CAN grid cell model created accurate path integration based on inputs that encode only a rat’s velocity using a firing rate coding (Burak and Fiete, 2009). Attractor states in the model occur as bumps of firing that are greater than areas outside of those regions. The study stipulates that grid cells have preferred directions, i.e., they are each specialized to respond more to certain direction-modulated input. An offset in the outgoing synaptic connections of grid cells influences directional movement of the bumps when activating those grid cells. A key property that helps generate bump activities is synaptic connectivity (and related weights) in center-surround (CS) structures. The CS structure supports bump firing in center regions and inhibits firing in surround regions. Another CAN grid cell model, using exponential integrate and fire neurons, included additional components such as place cells, signaling between grid cells and interneurons (IN), and three different synaptic receptor types (Solanka et al., 2015).

The present work uses neuron types and their properties from Hippocampome.org (Wheeler et al., 2015) to test whether previously proposed models can reproduce key physiological details observed in animal recordings when incorporating data-driven biophysical details. Hippocampome.org is a knowledge base of the rodent hippocampal formation circuitry and offers a wide variety of neural properties that are ready to be integrated into simulations (Wheeler et al., 2024). Parameters available from Hippocampome.org along with the corresponding experimental measurements include the Izhikevich Model (IM) of neuron firing and excitability (Komendantov et al., 2019; Venkadesh et al., 2019), Tsodyks-Markram (TM) model of synaptic signaling and short-term-plasticity (Beyeler et al., 2015; Moradi & Ascoli, 2019; Moradi et al., 2022), neuron counts (Attili et al., 2022), and connection probabilities (Tecuatl et al., 2021). Hippocampome.org also provides a wealth of neuron type-specific details about hippocampal activity and molecular expression that can add to the biological realism of computational simulations (Hamilton et al., 2017a; Hamilton et al., 2017b; Rees et al., 2016; Sanchez-Aguilera et al., 2021; White et al., 2020). We recently created a large-scale model of resting state dynamics in CA3 using Hippocampome.org properties (Kopsick et al., 2023) and showed its suitability to simulate pattern completion via cell assemblies (Kopsick et al., 2024). The present work complements those studies by also demonstrating the application of Hippocampome.org resources to modeling complex network interactions efficiently with reusable code.

This study’s software is released as open source to explore how varying property parameters within reported biological ranges affect the reproduction of electrophysiological data from mice. Comparing experimental recordings to simulated recordings assesses the ability of a physiologically realistic CAN model to recreate real neural activities. Testing of this model’s neural parameter predictions in rodents can verify their accuracy and be used to further refine the model.

## Methods

A recent review of publications on simulations of the hippocampal formation identified CAN grid cell models as a notable case relevant to spatial navigation (Sutton & Ascoli, 2021). Here, we used Hippocampome.org parameters to build a new SNN of grid cells adopting multiple methods from previous CAN models (Burak and Fiete, 2009; Solanka et al., 2015). In particular, we preserved the original model realism to only include inhibitory signaling between grid cells (an optional choice in Burak and Fiete, 2009) as opposed to other models that rely on excitatory signaling between grid cells (Spalla et al., 2019). Also, as in the original model (Burak and Fiete, 2009), preferred directions in grid cells managed bump movement, and each cell had a north, south, east, or west preferred direction. We adapted the model to convert its firing rate representation into the Izhikevich dynamical system formalism, which captures individual neuron spikes in a biologically realistic manner while also accommodating integration of Hippocampome.org parameters. We also used place cell input to help correct for drift in bump attractors as in a published CAN model based on exponential integrate-and-fire neurons (Solanka et al., 2015). We selected the TM synapse model using neuron type-specific parameters from Hippocampome.org for use in our work. In addition to factors similar to those included in previous models (e.g., Solanka et al., 2015), such as receptor-specific signal decay constants and reversal potentials, the TM model also includes variables for facilitation and depression, accounting for short-term plasticity.

### I. Mapping Functional Neuron Types to Morphological Neuron Types

Grid cells are most abundant in layer 2 (LII) of the medial entorhinal cortex (MEC), but also exist in other brain subregions such as the pre- and parasubiculum (Boccara et al., 2010; Sargolini et al., 2006). Grid cells in MEC LII can be either glutamatergic stellate cells (SC) or pyramidal cells (PC) (Kecskés et al., 2020). In rats, the primary population of excitatory cells in MEC LII are SCs (Dhillon and Jones, 2000; Witter et al., 2014). Those SCs express the large glycoprotein reelin (Kitamura et al., 2014; Miettinen et al., 2005) and largely influence each other through indirect connections via inhibitory interneurons. In contrast, PCs in LII may have direct connections with each other (Tecuatl et al., 2021). Given the different interaction mechanism and the empirical evidence on relative abundance, we chose to map grid cells to MEC LII SCs.

Anatomical, electrophysiological, and molecular information regarding GABAergic INs in the entorhinal cortex is notoriously scarce (Canto et al., 2008; Wheeler et al., 2024). Nevertheless, the identification of grid cells with MEC LII SCs provides definitive constraints as to the nature of the INs that could provide CS inhibition. In particular, because MEC LII SCs have soma and dendrites limited to layers 1 and 2, the INs that grid cells can use to communicate with each other must have axons extending into those layers. Moreover, parvalbumin-expressing (typically perisomatic-targeting) INs affect grid cell behaviors more than other IN neuron types (Miao et al., 2017). We thus chose to include in our model the three Hippocampome.org neuron types with those characteristics, namely LII axo-axonic, LII basket, and LII basket-multipolar cells, to capture a diversity of neural properties that may be present in animals (Fig. 1).

**Fig. 1.**
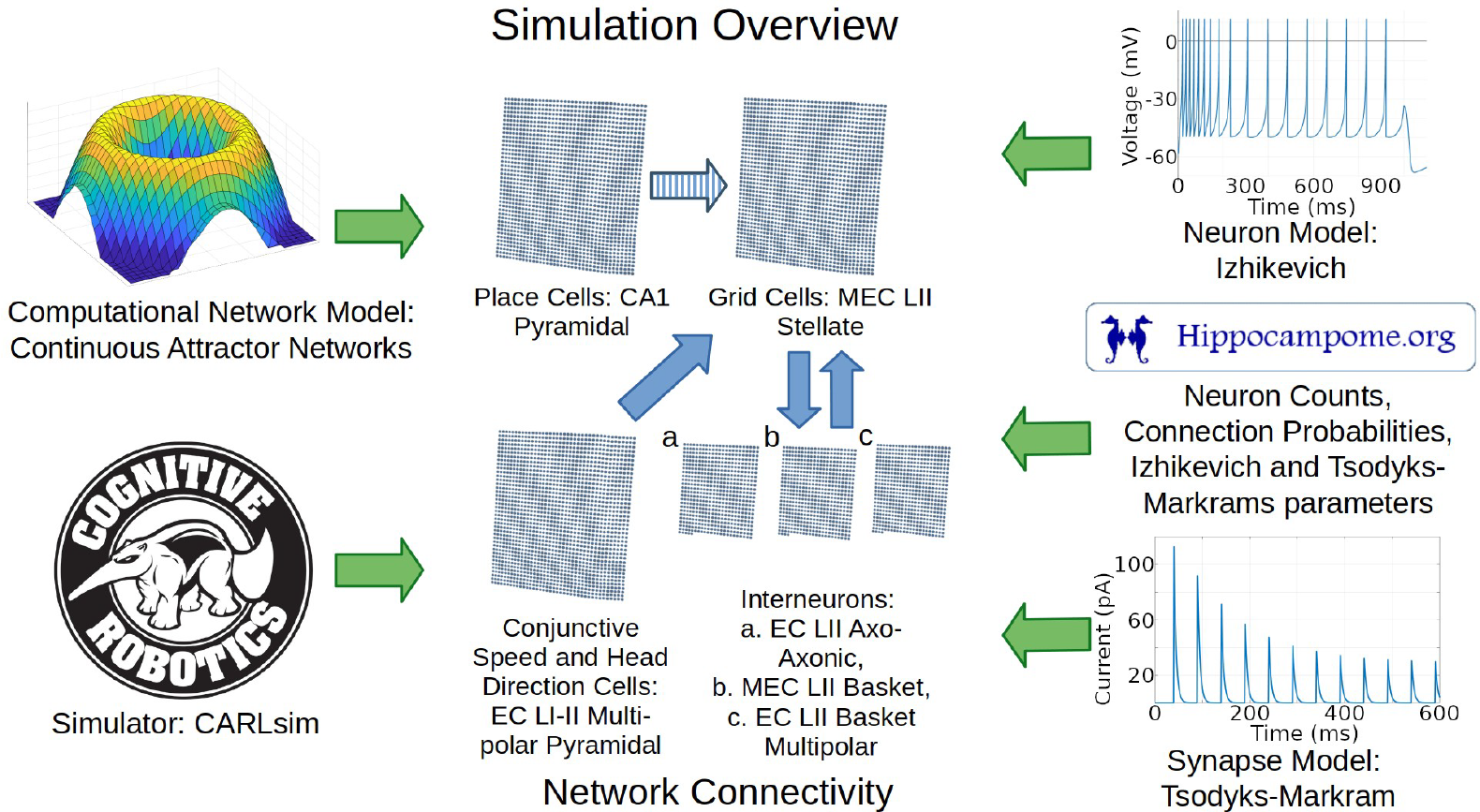
Overview of simulation components and methods. Green arrows represent components utilized in the simulation. Blue arrows represent directions of connections between the neural layers. The striped blue arrow indicates a connection between layers that may only exist indirectly in nature. Abbreviations: MEC, medial entorhinal cortex; LII, layer 2; CA1, cornu ammonis 1.

The neuron type chosen to represent place cells is CA1 PCs because those are well characterized neurons with place cell characteristics (Danjo, 2020). For simplicity, we simulated direct connections from place cells to grid cells. Indirect connectivity through deep layer MEC cells may be more realistic, though recent evidence also supports potential direct connections from CA1 to MEC LII (Butola et al., 2023). Our simplified abstract connection represents both the indirect pathway and the possible direct pathway, and is similar to the design adopted by prior models (Solanka et al., 2015). That similarity in network architecture helps test the effect of adopting data-driven parameters from Hippocampome.org while otherwise minimizing changes from previous methods.

Animal studies helped inform how to model the sources of signals that provide movement velocity information to grid cells. Coding for speed and direction contributing to velocity signals may come from separate neuron types or from conjunctive cells. One potential source of speed signal is from firing rates in a subpopulation of MEC neurons (Kropff et al., 2015). Head-direction cells (HDC) could provide running direction signals to grid cells, although HDCs can code more accurately for head direction than running direction (Raudies et al., 2015). HDCs have been identified in 11 brain regions including MEC LII (Dudchenko et al., 2019; Giocomo et al., 2011; Giocomo et al., 2014). Based on prior models and rodent studies, we included a cell group that provides velocity-modulated excitation to grid cells (Burak and Fiete, 2009; Solanka et al., 2015). We mapped these direction- and speed-modulated conjunctive cells to LI-II Multipolar PCs, the most abundant glutamatergic neurons in EC based on Hippocampome.org (Attili et al., 2022). These neurons have potential synaptic connections to stellate cells (Rowland et al., 2018; Tecuatl et al., 2021) and can thus supply direct excitatory input to grid cells (see Discussion for an alternative based on GABAergic signaling).

### II. Computational Modeling: Neurons and Synapses

We utilized CARLsim (Niedermeier et al., 2022) as the simulation platform because of its efficiency at high-performance computing of spiking neural networks using graphical processing units (GPU). The adoption of IM to describe neuronal excitability and firing allow simulations to recreate biologically realistic diversity and complexity of neuron type-specific input-output dynamics. This modeling formalism utilizes 4 parameters (K, a, b, and d) to control the intrinsic dynamics of the neuron, e.g., affecting bursting, rheobase, and spike frequency adaptation. Five additional IM parameters scale the neuronal response by setting the cell capacitance (C), resting voltage (Vr), threshold (Vt), spike cutoff value (Vpeak), and post-spike reset value (c). The 9 IM parameters (Table 1) were taken from Hippocampome.org for all glutamatergic neuron types. The available empirical evidence for MEC inhibitory cells is insufficient to constraint data-driven IM. Therefore, we used standard fast-spiking IM parameters suitable for these types of perisomatic, parvalbumin-expressing neurons (Izhikevich, 2007). The numbers of neurons in each type included in the simulation (Table 1) are also consistent with the Hippocampome.org cell census ranges for the mouse.

**Table 1.** IM parameters. ^a^ These values were utilized in the intermediate and small grid scale simulations (e.g., Fig. 2B-C). The number of INs was increased to 1200 in each of the three types for large grid scale simulations (Fig. 2A) to help accommodate control of the larger bump sizes.

| Neuron Type | Count | $C$ | $k$ | $V_r$ | $V_t$ | $a$ | $b$ | $V_{peak}$ | $c$ | $d$ |
| --- | --- | --- | --- | --- | --- | --- | --- | --- | --- | --- |
| MEC LII Stellate | 1600 | 118 | 0.62 | -58.53 | -43.52 | 0.005 | 11.69 | 11.48 | -49.52 | 0 |
| EC LI-II Multipolar Pyramidal | 1600 | 375 | 0.37 | -70.53 | -39.99 | 0.001 | 0.01 | 3.96 | -54.95 | 7 |
| CA1 Pyramidal | 1600 | 530 | 1.74 | -69.98 | -57.43 | 0.003 | -0.782 | 24.45 | -60.35 | 25 |
| EC LII Axo-Axonic | 834<sup>a</sup> | 20 | 1 | -55 | -40 | 0.15 | 8 | 25 | -55 | 200 |
| MEC LII Basket | 833<sup>a</sup> | 20 | 1 | -55 | -40 | 0.15 | 8 | 25 | -55 | 200 |
| EC LII Basket Multipolar | 833<sup>a</sup> | 20 | 1 | -55 | -40 | 0.15 | 8 | 25 | -55 | 200 |

We also strived to maximize the simulation biological realism in setting the properties of synaptic connections for each pair of neuron types, including the numbers of connections and the TM model parameters (Table 2). We considered both Hippocampome.org probabilities (Tecuatl et al., 2021) and additional animal data-driven values to select the connectivity ratios between neuron types. In particular, a study in mouse MEC LII reported 35.7% connectivity from INs to SCs and 25.9% from SCs to INs (Fuchs et al., 2016), while another mouse study reported a 26.9% connection probability in both directions (Fernandez et al., 2022). Additional studies have found higher connectivity between SCs and INs in rats (Couey et al., 2013; Grosser et al., 2021). The synaptic connections between INs and grid cells in the simulation follow a CS distribution of connection weights, with CS circumferences configured with the aim to match grid scales found in animal data, as detailed in supplemental material section (SMS) IV (the provided software helps automate construction of these distributions). All other connections had unitary weights and the model did not include long-term plasticity.

**Table 2.** Connection parameters. The values here are those used to produce the results in Fig. 2B. Values are rounded to the 3^rd^ decimal place. The number of connections column has numbers rounded to the nearest interval. ^a^ The connectivity was specific to each grid cell and varied within the standard deviation listed. ^b^ The individual number of connections reflect the CS distribution (details in SMS IV). ^c^ These values were adjusted for the scale of the simulation (details in SMS IV).

| Connection | # of Connections (mean ± SD) | $g$ (fast) | $\tau_d$ (fast; ms) | $U$ | $\tau_u$ | $\tau_x$ | $g$ (slow) |
| --- | --- | --- | --- | --- | --- | --- | --- |
| EC LI-II Multipolar<br>Pyramidal to MEC<br>LII Stellate | 1 to 1 ± 0 | 11.627<sup>c</sup> | 3.380 | 0.180 | 49.920 | 152.856 | 7.126<sup>c</sup> |
| MEC LII Stellate to<br>EC LII Axo-Axonic | 1 to 12 ± 2<sup>a</sup> | 1.050 | 3.085 | 0.167 | 49.920 | 167.686 | 0.644 |
| MEC LII Stellate to<br>MEC LII Basket | 1 to 12 ± 2<sup>a</sup> | 1.050 | 2.887 | 0.167 | 49.920 | 152.857 | 0.644 |
| MEC LII Stellate to<br>EC LII Basket<br>Multipolar | 1 to 12 ± 2<sup>a</sup> | 1.050 | 2.641 | 0.197 | 49.920 | 136.570 | 0.644 |
| EC LII Axo-Axonic<br>to MEC LII Stellate | 1 to 142 ± 66<sup>b</sup> | 0.626 | 4.561 | 0.145 | 27.306 | 420.838 | 0.383 |
| MEC LII Basket to<br>MEC LII Stellate | 1 to 142 ± 66<sup>b</sup> | 0.626 | 4.488 | 0.151 | 27.810 | 443.499 | 0.383 |
| EC LII Basket<br>Multipolar to MEC<br>LII Stellate | 1 to 142 ± 66<sup>b</sup> | 0.626 | 4.948 | 0.162 | 30.543 | 382.485 | 0.383 |
| CA1 Pyramidal to<br>MEC LII Stellate | 1 to 1 ± 0 | 9.248 | 3.380 | 0.123 | 49.710 | 153.400 | 5.668 |

The TM model calculates not only the intensity and duration of synaptic signals via a conductance (g) and kinetic decay (tau_d_) parameters, but also short-term plasticity with parameters (U, tau_u_, and tau_x_) controlling facilitation and depression (Moradi et al., 2022). Hippocampome.org provides data-driven ranges of TM parameters for fast (AMPA & GABA_A_) currents (Hippocampome.org/synapse). For each pair of neuron types, all TM parameters in our simulations were within two standard deviations of the Hippocampome.org means for 56-day old male mice at 32 degrees Celsius in voltage-clamp. This range represents 95% of the observed values under the assumption of a Gaussian distribution (Grafarend and Awange, 2012). We set the conductance level of slow currents (NMDA & GABA_B_) based on available evidence as described in SMS VII, and used the CarlSim default parameters for the NMDA and GABA_B_ time decays (tau_d_).

We created software programs to automatically test how simulations parameters within (and beyond) the ranges of Hippocampome.org IM neuron and TM synapse model values affected grid cell activity. Specifically, we adopted the fitness error from prior work (Venkadesh et al., 2019) to quantify the deviation of firing patterns obtained with a given selection of IM parameters from the Hippocampome.org experimental data for the same neuron type (Komendantov et al., 2019). Parameter exploration started from Hippocampome.org IM parameters and used an evolutionary algorithm (EA) software (as in Venkadesh et al., 2019) to test higher and lower values. We selected a firing pattern fitness error threshold of 200% of the minimum, which corresponded to the Hippocampome.org parameter value. This choice balanced accuracy and variability in reproducing the experimental firing patterns.

To evaluate how well the firing activity of a stellate cell reflected a grid pattern, we utilized a numerical score (grid score) previously introduced to quantify experimental data (Dannenberg et al., 2020). We chose a grid score threshold of 0.2 as an indicator of an acceptable grid pattern after a review of many plots with different scores. In these parameter exploration analyses, for computational practicality, we used a shorter simulation time than the full animal experiment time (see SMS I for additional details).

We analyzed spiking from the simulation to detect if rhythm-related properties were present. We created multi-taper power spectral density plots (to suppress noise) with Gaussian smoothing using a 3-ms moving window size (to minimize harmonic artifacts). We used a Rayleigh test to determine statistical significance in spike-phase coupling (Varga et al., 2012). We also computed mean resultant vector length (MRVL) values with the CircStat MATLAB toolbox (Berens, 2009) to quantify the strength of spike-to-phase coupling in the 0-1 range (Fernandez-Ruiz et al., 2023).

### III. Subjects and neural recording

We compared our SNN continuous attractor model of grid cells with benchmark experimental data from live behaving mice. Of 33 experimentally recorded grid cells utilized here, 28 came from a re-analysis of our earlier published work (Dannenberg et al., 2020), and 5 came from new recordings collected for this study. The earlier experiments investigated differences in grid cell firing between sessions recorded in light and darkness (Dannenberg et al., 2020). We only utilized the data from normal lighting conditions, i.e., the time periods that the lights were turned on.

The additional data were collected from 3 male mice (*Mus musculus*), 4 months old (1 wild type, C57BL/6J, and 2 heterozygous transgenic ChAT-Cre^tg/wt^. B6;129S6-Chat^tm1(cre)Lowl^/MwarJ; The Jackson Laboratory). All experimental procedures were performed in accordance with the regulations stated in the Guide for Care and Use of Laboratory Animals by the National Research Council and approved by the George Mason University Institutional Animal Care and Use Committee. Mice were implanted with a microdrive containing 4 movable tetrodes targeting the superficial layers of the left MEC at the following stereotaxic coordinates: 3.4 mm lateral to the midline and 0.5 mm anterior to the transverse sinus angled at a polar angle of six degrees. For an additional experimental purpose not reported in this study, a virus injection of calcium indicator GCaMP8s and a chronic implant of an optical fiber into the medial septum was performed. After surgery, mice were individually housed on a reversed 12 h light/dark cycle. The surgery for microdrive implantation and the electrophysiological recordings were performed as described previously (Dannenberg et al., 2020). Mice were given one week to recover from the surgery. Recordings of spikes and local field potentials were carried out while the animals randomly foraged for scattered pieces of cereal bits (Froot Loops, Kellogg Company, Battle Creek, MI, USA) in a wood open field environment of 45 × 45 cm^2^ with 30 cm high walls until animals sampled a sufficiently large area of the open field. One visual cue card (a white triangle) was placed on one of the walls of the open field. Tetrodes were turned in increments of 50 micrometers until spiking activity was detected and theta oscillations were observed indicating the tetrodes had entered the superficial layers of the MEC. Tracking of the position and head direction of the mouse was achieved by using head-mounted infrared LEDs. After data collection, animals were anesthetized deeply with isoflurane and transcranial perfusion with DPBS 1X + 10% buffered formalin was performed. To visualize tetrode track and MEC recording sites, sagittal brain sections (40-50 μm) were sliced using a vibratome and stained with a Cresyl violet solution.

The data analysis, including the classification of putative principal neurons, computation of spatial firing rate maps, and grid score, was performed as described previously (Dannenberg et al., 2020).

### IV. Hardware & Software

Results from this work were produced from running CARLsim6 on Linux with Ubuntu 21.10 and CentOS 8.5. We utilized both a stand-alone machine with a Nvidia GeForce RTX 3090 GPU and the supercomputer of George Mason University Office of Research Computing with a Nvidia A100 GPU. Typically, the simulation used 5 GB of RAM and 10 GB of GPU RAM. Additional software and hardware details are listed in the GitHub documentation.

## Results

Our research design consisted of utilizing experimentally recorded mouse locomotion to activate conjunctive (EC LI-II multipolar pyramidal) cells based on speed and direction, and place (CA1 pyramidal) cells based on position, as described further in SMS V. We then compared the corresponding simulated and real grid cell plots to gauge whether a spiking neural network implementation of the continuous attractor model with parameters based on experimental measurements could produce firing dynamics consistent with those observed in animal studies.

Simulations recreated well several properties of real mouse recordings, including grid field spacings, sizes, and orientation, as well as firing rates (Fig. 2). Multiple studies have reported grid field size and spacing increase from the dorsal to the ventral end of the MEC (Brun et al., 2008; Giocomo et al., 2014). This consistency of scale change along the dorsoventral axis suggests a purpose for this pattern. Our model could create grid fields with different scales qualitatively similar to those observed experimentally, such as large, intermediate, and small (Fig. 2A-C, respectively).

**Fig. 2.**
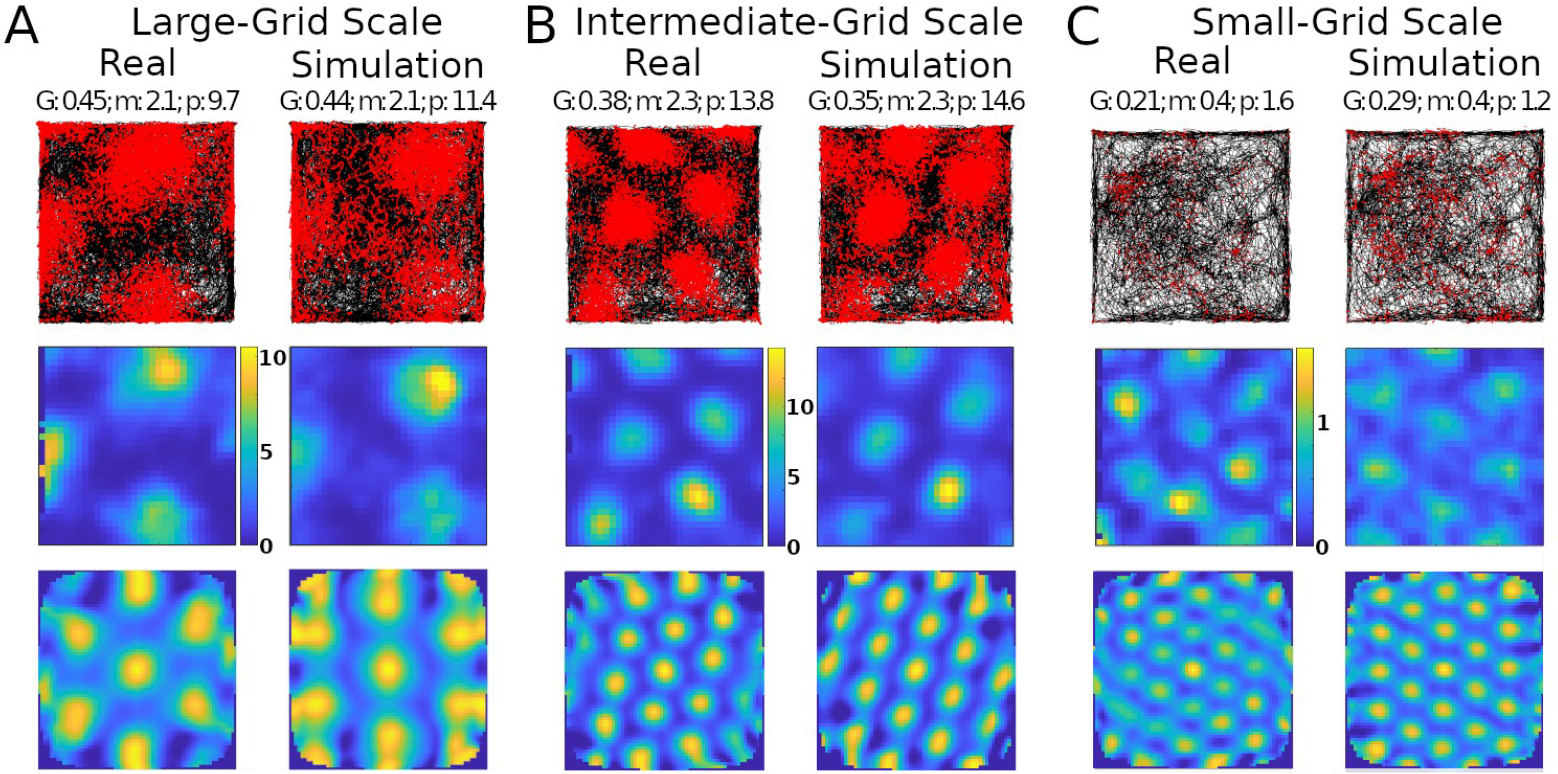
Comparing simulated and real results. Panels A, B, and C show results with different grid scales. Values reported at the top of the panels are grid score (G), mean firing rate in spikes/s (m), and peak firing rate, also in spikes/s (p). Top row: trajectory of animal in the environment (black lines) and spike location (red markers). Middle row: firing rate map of the neuron (color scale bars in spikes/s). Bottom row: autocorrelogram of the rate map (color bar ranges: A, -74% [percent is relative to maximum correlation] to 100%; B, -51% to 100%; C, -60% to 100%).

The simulations could generate specific rotations of grid patterns (see SMS V for methods) qualitatively similar to those observed in real recordings (Fig. 2A). When considering the whole analyzed time period, identical in real experiments and simulations (142.5, 141.4, and 60.2 minutes in Fig. 2A-C, respectively), the mean firing rate of the grid cells in Fig. 2 matched to a 1 decimal place accuracy. We thus compared the mean firing rates across an equal number of simulated and real grid cells (N=29) grid cells. For this purpose, we selected 6 simulated experiments to approximate the range of firing observed in real cells, including large-grid scales and low- and high-firing rates in small-grid scales. Scale sizes and firing rates were grouped based on the mean levels of the real cells (<= 50% for small or low and > 50% for large or high), and scale measurement methods are in SMS VI. In particular, the simulated small-scale cells included 11 low- and 9 high-mean-firing rate cells, and large-scale cells included 9 cells, which were similar to that observed in real cells. Because we could not assume normality of the mean firing rates, we opted for a non-parametric Wilcoxon rank sum test. The firing rates of the real cells had a median of 1.76 spikes/s and interquartile range (IQR) of 1.55 spikes/s, while the corresponding values for the simulated cells were 1.75 spikes/s (median) and 2.02 spikes/s (IQR) (Fig. 3). The p-value of 0.93 indicates a lack of support for the hypothesis that the firing rates of simulated and real cells have different distributions.

**Fig. 3.**
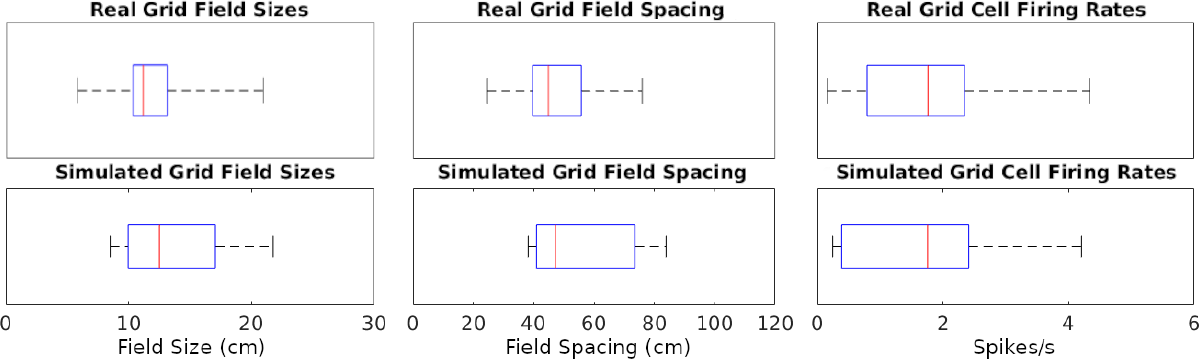
Distributions of grid cell metrics in real and simulated recordings. The data points analyzed are each cell’s mean of every value type (e.g., 7 field size values from a cell). Red lines are at median data points, blue rectangles represent data between the 25^th^ and 75^th^ percentiles, black whiskers extend to the first and last values in the sets of data points.

We also measured and compared grid field sizes and spacing between real and simulated recordings (see SMS VI for methods). These comparisons utilize the same cells used in the firing rates analyses and the Wilcoxon test for the same reasons described for firing rates. The grid field sizes of real cells had a median of 11.16 cm and IQR of 2.79 cm, and simulated cells had a median of 12.5 cm and IQR of 7.10 cm (Fig. 3). Grid field spacing in real cells had a median of 44.87 cm and IQR of 16.03 cm, and simulated cells had a median of 47.25 cm and IQR of 32.65 cm. The p-value for sizes was 0.41 and for spacing 0.26, both indicating a lack of support for the hypothesis that field sizes and spacings have different distributions between simulated and real cells.

Next, we investigated the robustness of the SNN implementation of the CAN model, as quantified by the grid score, relative to variations of neuronal and synaptic model parameters within their biologically realistic ranges. For IM models, focusing on the stellate cells, we considered parameter ranges to be biologically realistic if the resultant firing pattern had a fitness error up to 200% of the minimum, corresponding to the Hippocampome.org best ranking parameter value (see Methods for details). For TM models, we defined biological realism as two standard deviations from the mean of each parameter listed on Hippocampome.org. When testing the TM parameters, we focused on the synaptic connectivity from stellate cells to INs, varying the values of all three sets of glutamatergic connections (onto axo-axonic, basket, and basket multipolar cells) at the same time.

This analysis demonstrated that the emergence of grid score dynamics in the CAN model is more sensitive on certain model parameters than others (Fig. 4 and S3). In particular, several IM and TM parameters have wide ranges of variability that generated above-threshold grid scores. These include stellate cell IM parameters k, a (Fig. 4A), V_t_, V_peak_ (Fig. S3B), and V_r_ (Fig. S3C), as well as stellate-to-IN TM parameters g, tau_d_ (Fig. 4B), and tau_u_ (Fig. 4D). This indicates that grid field functionality could be preserved amid a broad diversity of physiological modulatory factors that could affect the intrinsic excitability and synaptic signaling of MEC LII stellate cells. In contrast, the range of viable values for TM parameter tau_x_ producing sufficient grid scores was substantially more limited (Fig. 4D). This indicates that tau_x_ may be specialized toward accommodating grid field activities. The situation is more complex for IM parameters b and d. While each of them has fairly broad ranges of values yielding strong grid patterns, they also depend on each other via negative correlation: the highest values of parameter b only allow high grid scores when paired with the lowest values of parameter d, and vice versa (Fig. 4C). A similar relation is present between IM parameters C and V_min_ (Fig. S3A). The areas within thresholds in these plots help reveal the sensitivity of each score to the parameters.

**Fig. 4.**
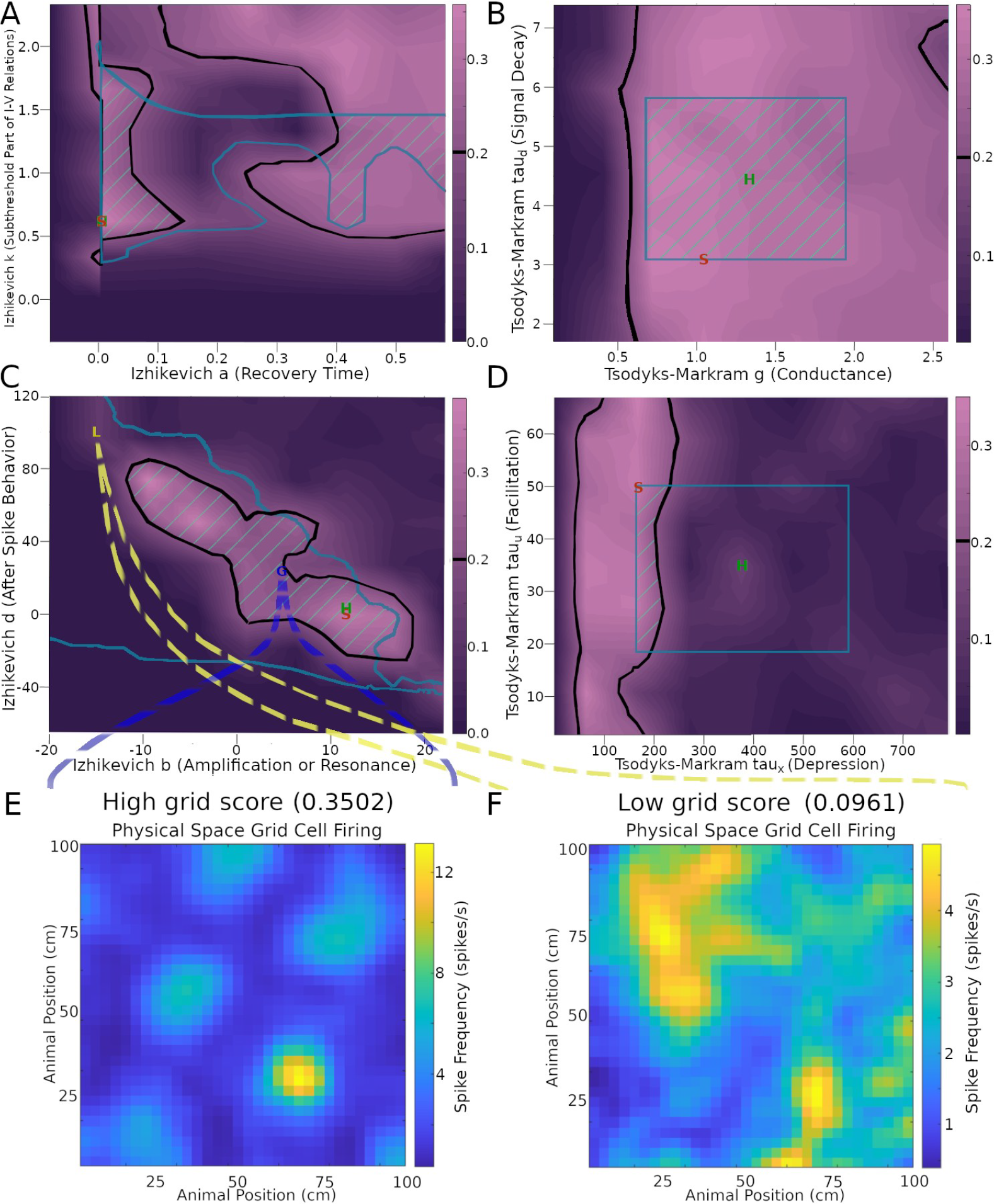
Effects of neuronal and synaptic model parameters on stellate cell grid scores. The plots in panels A-D were obtained by testing nine values per parameter and interpolating scores between those values. Black outlines encompass regions with grid scores above the 0.2 threshold (cf. color scale bars on the right of the plots). Teal outlines indicate the biologically realistic parameter ranges. Green stripes represent intersection areas with both acceptable grid score and biologically realistic model parameters. Green ‘H’ and red ‘S’ letters mark the default parameter values on Hippocampome.org and the values used in the simulation for Fig. 2B, respectively. Panels E and F plot the firing rate maps obtained with representative settings inside and outside the acceptable parameter spaces (G and L markers, respectively, in panel C).

To illustrate the effect of these biophysical parameters on grid cell activity, it is useful to examine examples of the firing rate maps. In particular, values from areas of the parameter space giving rise to a high grid score yield strong grid patterns (Fig. 3E), while values outside of those areas cause the firing to lose its grid pattern (Fig. 3F). Interestingly, the average firing rate remains approximately the same in both cases (∼ 2.5 spikes/s in panel E and F). However, the firing activity is largely focused onto field areas in the high-grid score case (Fig. 3E) while it is widespread in the environment in the low-grid score case (Fig. 3F). Of note, the simulation settings used to produce the grid scores and other results in Fig. 3, aside from parameters explored, remained constant and were the same as those used in Fig. 2B, corresponding to an intermediate grid scale cell. The results remained reasonably consistent when extending the parameter search analysis to small- and large-field cells (Fig. S4).

One of the advantages of a SNN implementation of the CAN grid cell model is the ability to inspect firing dynamics from single cells to the entire circuit. Throughout the simulations, network activity yielded realistic, sustained, non-repetitive spiking patterns (Fig. 5 and Supplementary Video at Hippocampome.org/RTPlots2). Specific activity patterns emerged in each collection of neuron types. At any given time, only a subset of grid cells is active, and the constellation of this subset changes smoothly over time generating the appearance of curved lines in the raster plot, consistent with an activity bump moving along a continuous attractor space. Stellate cell firing stimulated INs, increasing their spiking immediately after the activation of grid cells. In turn, INs inhibited the grid cell layer in different locations due to the CS connectivity. Conjunctive cell spiking reflects a combination of a baseline firing level and activity targeting grid cells with specific preferred directions. This results in nearly all conjunctive cells firing together at times with only specific groups of cells spiking in between bouts of synchronous activity.

**Fig. 5.**
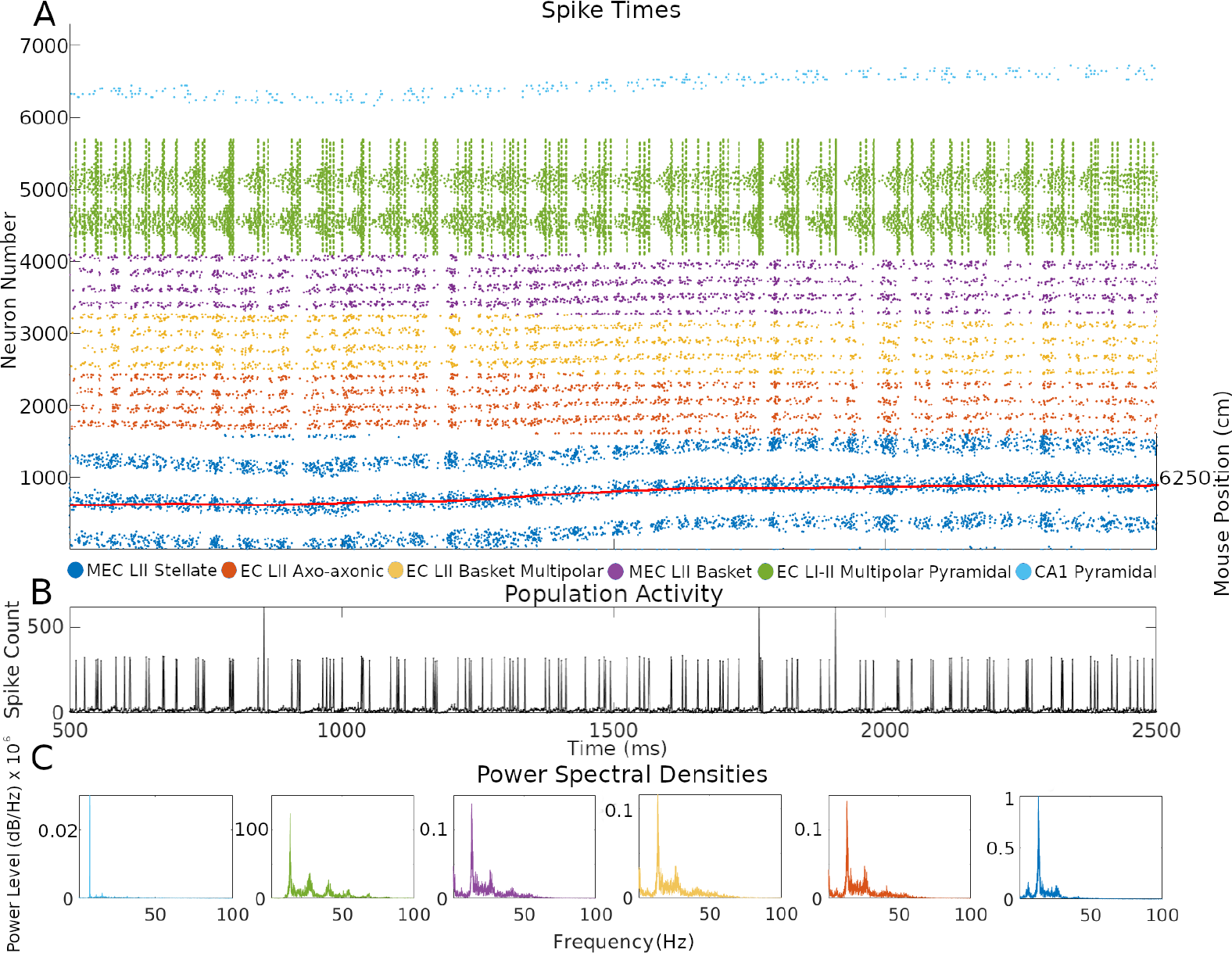
Neuron type-specific spiking dynamics. (A) Raster plot of the spiking of all neurons during two seconds of simulation of animal data. This activity was from the simulation in Fig. 2B. The red line represents the movement path of a mouse that was used as velocity input into the simulation: the corresponding right y-axis label represents two-dimensional environment position converted to one-dimension with the formula (*y* – 1) * *x*_*size*_ + *x*, where y is the y-axis location position, x is the x-axis location position, and *x*_*size*_ is the length of the x-axis in cm. (B) Population activity from the combined spiking of all simulated neurons in MEC LII, namely grid cells, INs, and CjCs. (C) Power spectral densities of neural spiking during the first minute of the simulation. The spiking in each plot is from all neurons within each color-coded neuron type group.

CA1 PCs showed relatively sparser firing compared to other neuron types. This is consistent with the known physiology of these neurons and with the function of place cells, which only respond to restricted areas of an environment. The single bump of CA1 PC place fields limits their firing to a relatively narrow group of neurons during the course of a few seconds. In contrast, grid cells, and the INs connected to them, fired in multiple places due to the multi-bump structure of the CAN. The intended movement velocity of the animal also affects the level of firing at each time. A speed control function differentially activates conjunctive cells depending on the speed of the animal, and that signal propagates throughout connected cells (SMS V). Conjunctive cells also more strongly stimulate grid cells with preferred directions that match the direction of animal movement at each timestep, which contributes to signaling changes.

Next, we investigated the rhythms of the network activity displayed in Fig. 5 by power spectrum density analysis using the combined spiking activity of each neuron type group. When considered all together, MEC LII neurons had the highest power in the beta band (12-25 Hz), followed by the gamma band (25-100 Hz), with 41.7% and 25.5% of total power, respectively. Taken individually, Stellate, Axo-axonic, Basket, and Basket Multipolar cells had their highest powers in the beta band (55.9%-44.6%), and second highest in the gamma band (33.4%-24.8%; Fig. 5D). In the hippocampal formation, phase coupling is especially informative relative to the slow network oscillations (Klausberger & Somogyi, 2008; Kopsick et al., 2023; Sanchez-Aguilera, 2021). We selected the peak firing of stellate cells within the beta rhythm as a common frequency to evaluate MRVL values of all neuron types during the first minute of the simulation. We based this choice on the stellate cells being directly connected to all other neuron types and the beta rhythm having the highest power across neuron types (Fig. 5C). Stellate, axo-axonic, basket, and basket multipolar cells had a MRVL value of 0.04, and multipolar pyramidal cells had a MRVL value of 0.05. These neuron types had Rayleigh test p-values smaller than 0.001. These results indicate these neuron types had significant but weak spike-phase coupling to the beta rhythmicity in grid cell firing (Bezaire et al., 2016).

The activity of the grid cells in their corresponding neural space reflected the multi-bump structure of the CAN model (Fig. 6A and Supplementary Video). At the same time, the spiking neural network implementation of the CAN model also allows the virtual recording of voltage and current in a selection of its simulated neurons (Fig. 6B-D and Supplementary Video). The current plots recorded in the stellate cells combine the contributions from both fast (AMPA and GABA_A_) and slow (NMDA and GABA_B_) receptors from all sources, namely CA1 pyramidal, EC LI-II multipolar pyramidal, EC LII axo-axonic, MEC LII basket, and ECLII basket multipolar cells. The currents recorded in the axo-axonic cells correspond to the excitatory input (AMPA and NMDA) from stellate cells.

**Fig. 6.**
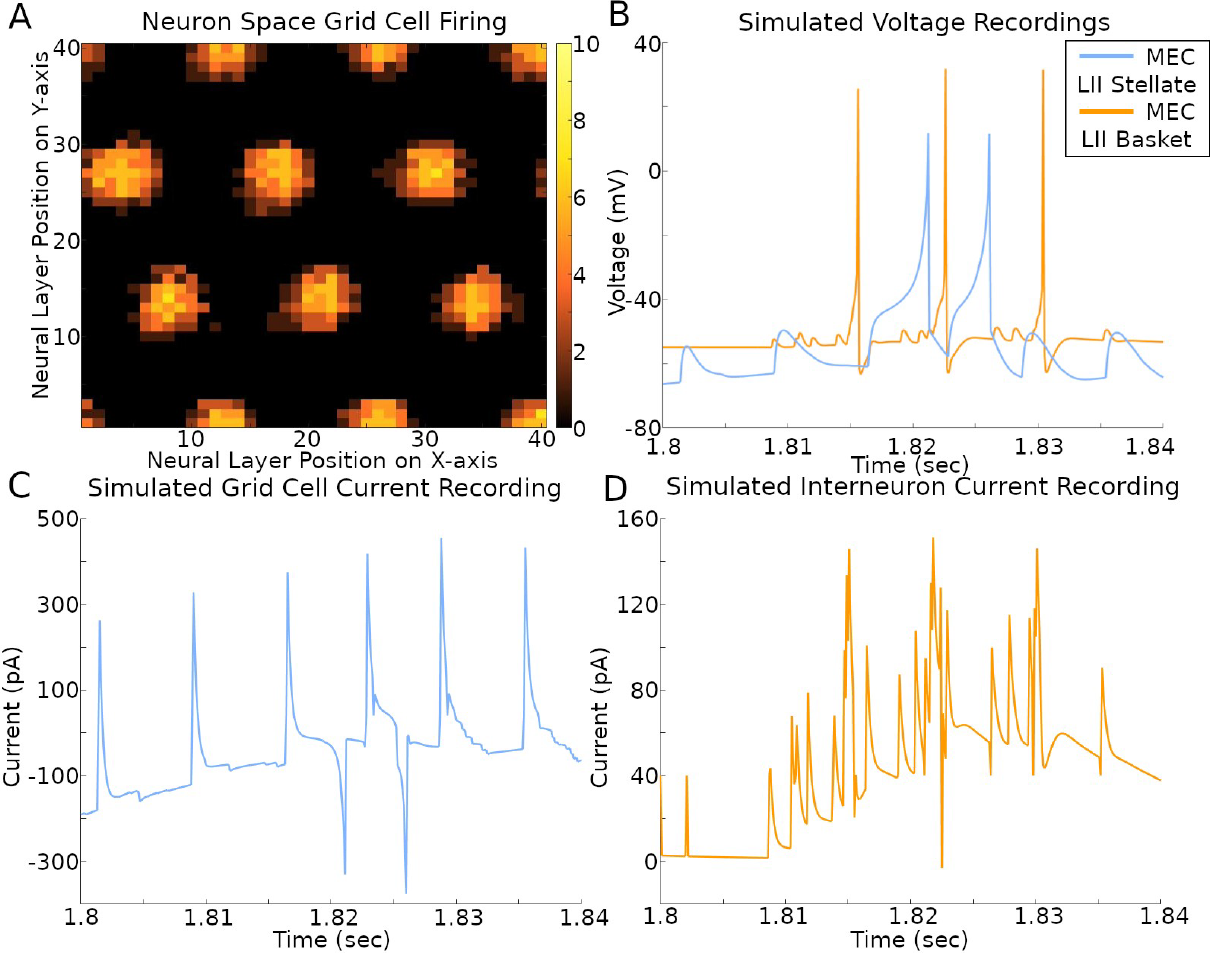
Simulated voltage and current. The simulation settings were those used to create results in Fig. 2B. (A) Rate map of all grid cell spiking within a 400 ms time span. The color bar on the right is the count of spikes in that time period. (B-D) Recordings during that same time span of a stellate cell and IN with x- and y-axis position (2, 1) in their respective neural layers.

In order to facilitate future related research, we released the simulation software pertaining to this work as open source at Github.com/Hippocampome-Org/spatial_nav. We also release as open access all mouse recording data from (Dannenberg et al., 2020) and 5 new cells recorded for this work at (Sutton et al., 2024a; Sutton et al., 2024b). Among several functions, the released software automates parameter exploration and generates recording scores and plots. Researchers can use this free resource to reproduce the simulation results, further investigate the spiking network model, expand on this work, and repurpose parts or the entirety of the code for studies of different subjects. The software includes usage instructions and documentation on its components, covering a wide range of topics to guide users from beginner to advanced levels of work with these simulations. Additional details on hardware and software versions used in this study are also available in the documentation.

## Discussion

Using a biophysically detailed simulation, we recreated well several properties of grid cells observed in animals, including field spacing and size as well as rotation of grid patterns. Furthermore, the simulations recreated the increase in the spacing and size of individual grid fields similar to those observed in rodents along the dorso-ventral axis of the MEC (Hafting et al., 2005; Pastoll et al., 2012). Grid cells can be grouped into discretized modules that share similar field sizes and spacings, grid pattern orientation, and other characteristics (DiTullio and Balasubramanian, 2021; Moser et al., 2014; Stensola et al., 2012). These modules have been theorized to represent different spatial scales of an environment in a computationally efficient manner (Wei et al., 2015; Mosheiff et al., 2017; R.G. et al., 2024) and could work together in spatial cognition to inform an animal of its location (Burak, 2014; Waniek, 2020). Controlling field spacing and size in the simulation helped recreate properties of different grid cell modules throughout a realistically broad range of scales. This opens the prospect of utilizing this spiking neural network implementation of the CAN in future explorations of the computational function of multi-scale grid representation.

Our simulation also reproduced with good accuracy several characteristics of the grid cell firing rates observed in real animals, including the average, the variability, and the differences between in-fields and between-field levels. Prior CAN model implementations typically created homogeneous firing fields, contrasting the natural heterogeneity of field firing and shape in experimental recordings. The firing variability of our simulation was an emergent property of the data-driven neural and synaptic properties modeled based on empirical evidence sourced from Hippocampome.org. Given the potential importance of variable neuronal firing in network function (Stein et al., 2005), it may be interesting to investigate computationally the role of heterogeneous grid cell activity in spatial navigation and pathfinding.

We deemed it valuable to recreate grid pattern rotations relative to the environment borders because of studies indicating that those rotations could be non-random and functionally relevant. In particular, the offset angle of the grid pattern axes from the walls minimized symmetry with the area borders (Stensola et al., 2015). Moreover, moving a cue-card along the wall of a circular environment coincided with grid cell pattern rotations (Hafting et al., 2005; Solstad et al., 2008). Interestingly, rotation of geometrically polarized enclosures (e.g., square walled areas) affected grid pattern rotation more than distal cues (e.g., rectangular cue card), but in geometrically symmetrical enclosures (e.g., circular areas) the rotation of distal cues had greater effects on grid pattern orientations (Krupic et al., 2015).

The data-driven SNN implementation of the CAN released with this work could be useful in the future to investigate the biophysical and circuit mechanisms underlying the geometric relationship between the spatial representation of environmental stimuli and grid patterns.

Parameter exploration identified ranges of biophysical values supporting grid cell functionality, and those ranges constitute testable hypothesis bridging quantitative physiology and circuit function. For instance, future studies could attempt to fit the firing patterns of putative MEC LII SCs recorded in vivo with IM models. This would allow a statistical comparison of the distributions of IM parameters corresponding to grid and non-grid cells against the predictions of our simulations. In principle similar experiments could be designed to fit synaptic activity between grid cell and INs with TM model parameters, although in vivo recording of unitary synaptic signals remains technically challenging in live behaving animals.

The synaptic connectivity in the simulation was constrained by values reported for each directional pair of neuron types by Hippocampome.org and the peer-reviewed literature. For each presynaptic and postsynaptic neuron type, Hippocampome.org estimates the overall probability of connection for the whole MEC based on axonal and dendritic morphology. In contrast, most experimental studies quantifying connectivity, typically based on paired neural recordings, rely on a limited sample of neurons in relative local proximity to each other. We designed our simulation with an intermediate connectivity between these two extremes, consistent with the modeled neuron counts in each type falling between full and local scale. For instance, because experimental studies reported 25.9% connectivity between SCs (potential grid cells) and INs in mice, we constrained the connectivity in our model so as not to exceed that amount. The effective simulation of cells with biologically realistic grid patterns and high grid scores using this connectivity indicates potential matches to real cell properties, but more animal recordings are needed to fully characterize circuit connectivity.

In rats, more connections are present between LII basket cells and principal cells in the dorsal region as compared to the ventral region of the MEC (Grosser, 2021), including SCs that could be grid cells. The CAN model in this work followed a similar pattern of connectivity with grid cell to IN connections relative to grid scales. When simulating large-grid scales, found in ventral MEC, only a limited number of inhibitory CS rings fit within the neural layer of grid cells before notably overlapping with each other, which caused unwanted bump movements. In our SNN simulation, this CS inhibition is generated by fast-spiking INs, which are activated by excitatory synaptic connections from grid cells. Smaller grid scales, found in dorsal MEC, allowed many more rings without overlap, and therefore a greater number of grid cell to IN connections with effective bump movements. The approximate average number of 5, 12, and 16 IN connections from each grid cell, in large-, intermediate-, and small-grid scale simulations, respectively, follows the same connectivity trend reported along the dorsoventral axis. A more in-depth analysis of circuit connectivity in our simulation is available in SMS IV.

Future rodent recording can test if similar features of the firing patterns observed in the simulated raster plots also emerge in vivo. Since the simulation only includes a subset of MEC neuron types, the comparison would require techniques to distinguish those kinds of neurons, such as stellate cells, while recording thousands at a time with a 1 millisecond temporal resolution. When available, we predict that such experiments will reveal grid cell activity occurring in curved line patterns as observed in our simulated raster plots, given that the cell firing reflects animal movement. Such firing is predicted to occur along a toroidal manifold (Gardner et al., 2022). To display that pattern, it will be sufficient to arrange the grid cells in the raster plot in sequential order of their response to the animal positions in the environment. Moreover, the periodicity of grid cells emerging from a sequence code of trajectories (R.G. et al., 2024; Zutshi et al., 2017) in an environment could produce various patterns of how the activity bump moves in time, where the pattern is informative of the trajectory of the animal, as observed in our data. The firing of grid cells in the simulation within activity bumps often follows closely the movement of the mouse (Fig. 5B).

One of the brain structures targeted earliest by Alzheimer’s disease is the hippocampal formation, including the entorhinal cortex (Gail Canter et al., 2019; Schmitz & Nathan Spreng, 2016), as characterized by altered neural excitability (Angulo et al., 2017) and synaptic dysfunction (Sheng et al., 2012). Spiking neural network simulations exploring how such perturbations could affect grid fields, in parallel with real recordings in animals engaged in a spatial task, could help bridge the gap between data from animal experiments and human relevance, and lead to early-detection or treatment advancements. In addition to investigating disease, biologically realistic computational models such as developed in this work could more generally foster a deeper understanding of the role of attractor dynamics in multiple forms of learning and memory (Spalla et al., 2021).

In our simulations, similar to prior CAN models, a single layer of excitatory neurons provides direction and speed input signals to grid cells. However, speed signals could also come from inhibitory INs (Ye et al., 2018), and a near real-time accurate speed code by firing rate may require fast-spiking INs (Góis and Tort, 2018; Dannenberg et al., 2019). Another alternative or complementary source of speed signal could come from the correlation of speed with theta rhythms (Hinman et al., 2016; Dannenberg et al., 2019; Dannenberg et al., 2020) and related cholinergic (Dannenberg et al., 2016; Kopsick et al., 2023) or glutamatergic (Fuhrmann et al., 2015) input from the medial septum. Expanding this SNN implementation of the CAN to include those additional mechanisms might help shed light on their computational plausibility.

In the simulation, grid cells rely on place cell input to assist with drift correction. Boundary cells, visual landmarks, and other mechanisms may also contribute to drift correction (Stratton et al., 2011). The place cell input used in this simulation causes extra firing to appear in one grid field relative to others. While this is often observed in real animal data (Dannenberg et al., 2020), the current model might have challenges in exactly reproducing the activity of grid cells with uniform firing fields. Well-designed automated speed control functions in this model can help reduce the need for drift correction, but incorporating additional drift correction methods might aid in capturing more types of grid field firing. Tightly coordinated computational and experimental research could reveal major insights into the mechanisms of drift correction.

In rare instances (<2% of recorded time), the animals traveled especially fast, e.g., 90 cm/s. We considered those speeds as outliers and did not tune the model to match those cases. Attempting to optimize the simulations for outlier speeds with the current design decreased the accuracy of the results at slower speeds. This was the case when an animal rapidly transitioned from extremely fast movements to very slow speeds, causing a spurious neural signal to linger during slower speeds. It would be interesting for future research to explore the circuit dynamics underlying these transitions in parallel animal experiments and computational simulations.

## Conclusion

This work integrated detailed neural and synaptic properties from Hippocampome.org and the experimental literature to enhance the realism of a grid cell CAN model. The results showed many qualitative and quantitative similarities to real recordings. Exploration of the effects of multiple physiological parameters on grid patterns narrowed down the biologically realistic ranges that may enable proper spatial navigation, generating testable hypotheses for future animal studies. Reproduction of key animal recording features with good accuracy, while integrating neuron type-specific empirical data on cell excitability and synapse signaling, lends support for the CAN model of grid cell activity.

## Supplementary Materials

The supporting information can be downloaded at www.mdpi.com/example/link (to be added once the link is available).

## Author Contributions

N.S. performed the simulated experiments. B.G. carried out animal experiments. G.A. and H.D. supervised the project. G.A., N.S., and H.D. designed the simulation experiments. H.D. and B.G. designed the animal experiments. N.S. and G.A. drafted, and H.D. and B.G. revised, the paper. All authors have read and agreed to the published version of the manuscript.

## Institutional Review Board Statement

The study was conducted according to the guide for the care and use of laboratory animals published by the National Research Council and the Institute for Laboratory Animal Research in the United States. All experiments were reviewed and approved by the Institutional Animal Care and Use Committee of George Mason University.

## Data Availability Statement

Data of this study are available from online links described in this manuscript. Any additional data are available from the corresponding author upon reasonable request.

## Informed Consent Statement

Not applicable.

## Acknowledgments

The authors thank the following people for sharing their knowledge and generous support toward this project: Dr. Diek Wheeler, Jeffrey Kopsick, Dr. Keivan Moradi, Dr. Siva Venkadesh, Dr. Ketan Mehta, Dr. Bengt Ljungquist, Dr. Carolina Tecuatl, Dr. Jeffrey Krichmar, Lars Niedermeier, and all others who helped provide ideas.

