## Supplemental Document for "A Continuous Attractor Model with Realistic Neural and Synaptic Properties Quantitatively Reproduces Grid Cell Physiology"

I. Additional parameter exploration methodological details

The real animal trajectories used in our modeling setup extend over a substantial time span. For instance, the trajectory utilized in the simulation of Fig. 2B, displayed in Fig. S1A, took 141.4 minutes. Since parameter exploration (whose results are reported in Fig. 3) entails a large number of simulations, in those cases we increased computational efficiency by adopting an alternative movement trajectory, shown in Fig S1B, that covered much of the same 10000 $cm^2$ area (2025 $cm^2$ in some cases) in ~4.2 minutes. The online documentation provides additional details about work with the trajectories.

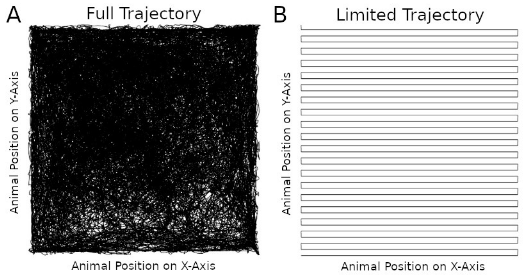


Fig. S1. Trajectories of movement used in recreating Fig. 2B's animal recordings (A) and parameter exploration (B).

Utilizing the limited trajectory instead of the full trajectory still allowed a reasonably accurate reproduction of the grid cell rate maps recorded from real animals (Fig. S2). Additional details on parameter exploration settings are in the online documentation.

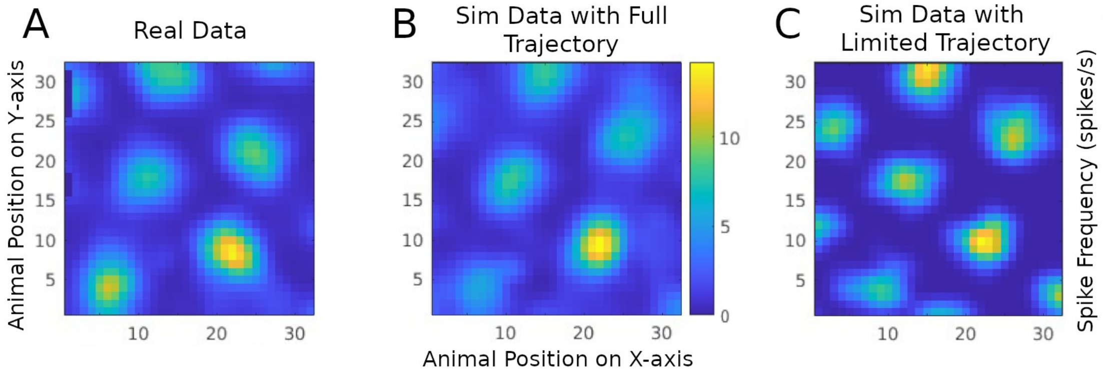


Fig. S2. Comparison of rate maps from a real animal recording (A), a simulation utilizing the full trajectory (B), and simulation utilizing the limited trajectory (C).

The parameter exploration settings generated a grid score of 0.36 before any parameters were modified.

The grid score threshold of 0.2 was selected after reviewing plots with permuted real animal data: starting with a high-scoring real animal plot, an automated script incrementally permuted the firing times of spikes to degrade the grid pattern, lowering the score.

II. Additional parameter exploration results

To select parameter ranges for testing (Fig. S3) we started from the original ranges included in the evolutionary algorithm (EA) software settings for parameter searching (Venkadesh et al., 2019). For Izhikevich parameters C and $V_t$, we used similar ranges as in the software. For Izhikevich parameters $V_{min}$, $V_{peak}$, and $V_r$, we expanded the ranges, as the original settings only spanned 1 unit each. The results indicate that the grid score remains robustly above threshold for broad ranges of parameters $V_{min}$ (Fig. S3A), $V_t$ and $V_{peak}$ (Fig. S3B), $V_r$ (Fig. S3C) and $Tau_u$ (Fig S3D). In contrast, the grid score is more sensitive relative to parameters C (Fig. S3A), d (Fig. S3C), and U (Fig. S3D).

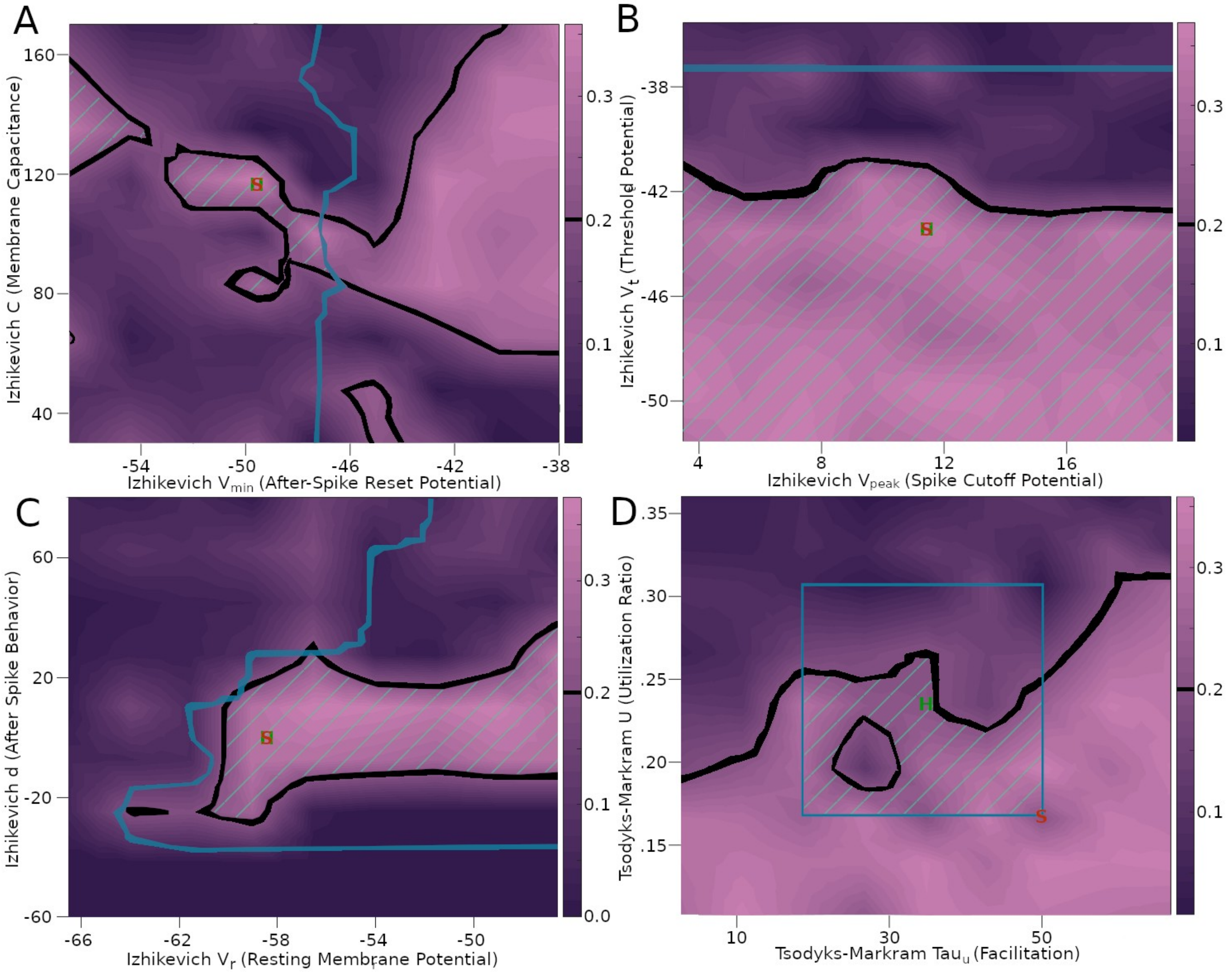


Fig. S3. Parameter exploration with Izhikevich and Tsodyks-Markram parameters. Each panel includes a different combination of parameters tested. The colors and highlighted areas of the images are the same as explained in Fig. 3C.

We also extended the parameter exploration to simulations recreating different grid scales (Fig. S4). The results confirmed the existence of many combinations of IM parameters that produce grid scores above the 0.2 score threshold. This indicates that grid cells may be produced by a variety of parameter values at different grid scales.

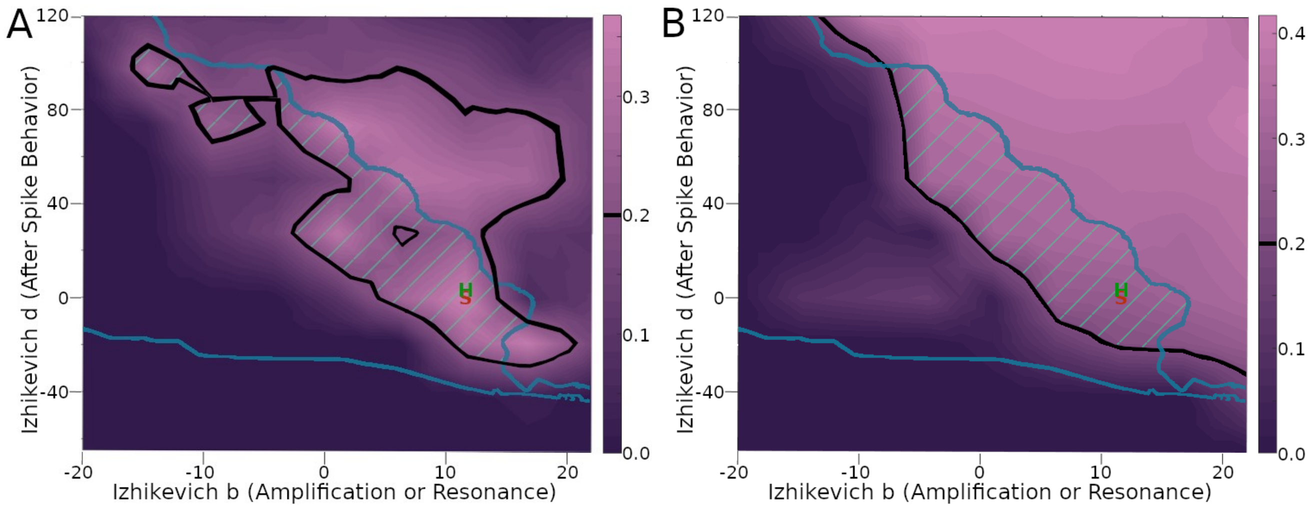


Fig. S4. Parameter exploration with grid cells that have different grid scales. (A) Izhikevich model (IM) parameter variations with large-grid scale cell (cf. Fig. 2A). (B) IM parameter variations with small-grid scale cell (cf. Fig. 2C). The colors, “H” and “S” markers, and highlighted areas of the images are the same as explained in Fig. 3C.

III. Animation of signals occurring in real time

A video from a simulation with the settings from Fig. 2B is available at Hippocampome.org/RTPlots2 (screenshot in Fig. S5). The video covers approximately the first 5 minutes of the simulation followed by another 5 minutes that occur 1 hour after the start of the simulation. The spike raster includes the first 50 neurons in the grid cell layer. The neuron locations are identified with the position formula described in Fig. 5’s caption. We post-processed the voltage plot to extend the spike upstroke to 40 mV when the membrane potential exceeded 0 mV. This adjustment yields a more interpretable visual rendering of action potentials given the limitation of the CARLsim simulation software to store in a file the peak voltage at the exact spiking time for plotting.

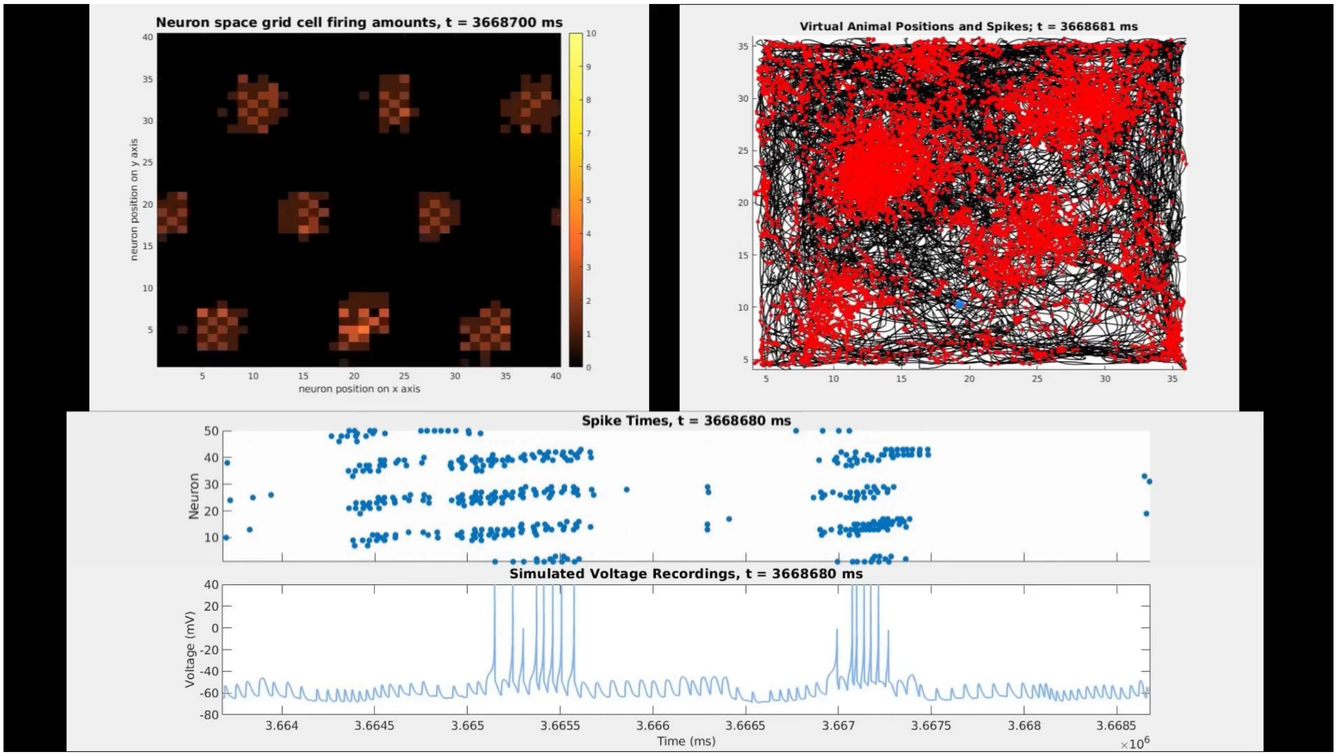


Fig. S5. Screenshot of video with recordings plotted in real time. Top left: rate map plot in the same style described in Fig. 5A. Top right: animal trajectory plot as described in the top row of Fig. 2. This trajectory plot and the neuron's membrane potential shown in the bottom panel are from neuron 1 in the grid cell layer. The middle panel is a raster plot of the same type described in Fig. 4.

### IV. Connectivity modeling

#### a. Connectivity design and statistics

We selected the simulation connectivity parameters using two complementary data sources: electrophysiological studies and Hippocampome.org. Electrophysiological studies based on either paired recording with intracellular electrodes in slices (e.g., Fuchs et al., 2016) or spike-time cross-correlograms from tetrode recording in vivo (Buetfering et al., 2014) often report connectivity between cells that have somata close together in 3D space. Grid cells may also connect with other neurons whose somata are farther than typically probed with paired recordings. Hippocampome.org represents the full-scale circuitry of the hippocampal formation, including long-range projections, and thus reports connectivity that is usually lower than that estimated in paired recordings. Our simulation has an intermediate neuron count between the whole medial entorhinal cortex and the local circuit accessible to electrophysiological recordings. Thus, we used the connectivity values from Hippocampome.org and electrophysiological recording studies as lower- and upper-bound constraints on connectivity for the model.

Of note, in our simulation each grid cell connects to a few interneurons (INs), which causes more than one center-surround (CS) inhibitory distribution to be signaled back to grid cells from INs. This approach differs from prior models, in which each grid cell enabled signaling from a single CS distribution to other grid cells, corresponding to a one-to-one grid cell to IN connectivity, as opposed to the one-to-few design adopted here. Including these multiple grid cell to IN connections also helps protect against a bump drifting error. Bump activity takes the form of a grid pattern (in neuron space firing rate map plots, as illustrated in Fig. 5A), which largely influences the grid pattern recorded in the firing of individual neurons (in physical space cell firing rate map plots, as illustrated in the second row of Fig. 2). Such grid patterns in neuron and physical spaces can be rotated around the bump/field at the center of the pattern. The drift error observed is that these grid patterns can drift into different rotations during the simulation. To achieve grid field firing in physical space plots that is similar to those found in animal data, it is important for the rotation to remain consistent instead of changing. The multiple grid cell to IN connections produce firing at a specific rotation rather than a mixture of one or more of them resulting from a one-to-one connectivity. Additional design details related to the connections are in documentation at Github.com/Hippocampome-Org/spatial_nav.

We chose approximately equal population sizes for the three IN types (compatible with the statistical distributions reported by hippocampome.org/php/counts.php) to ensure a diversity of GABAergic signaling in the simulation. Each IN cell inhibits grid cells in a center-surround ring pattern using a staggered formation to intermix the IN effects. Specifically, we alternated the assignment of CS rings between the first, second, and third IN type sequentially across adjacent grid cells (Fig. S6). Future studies could investigate whether this connectivity, or variations thereof, may occur in real animals.

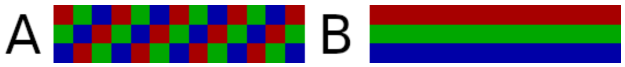


Fig. S6. Schematics of center-surround connections from the three IN types to the grid cells, where each square represents a targeted grid cell ring center, and the colors correspond to the distinct IN types. (A) Staggered pattern used in our simulation. (B) Possible non-staggered alternative shown for comparison.

A recent pair recording study of the inhibitory feedback microcircuits between principal excitatory cells and basket cells in rat brain slices found a higher-to-lower connectivity gradient along the dorsal-to-ventral axis of layer 2/3 in the medial entorhinal cortex (Grosser et al., 2021). The dorsal and ventral regions generally have smaller and larger grid fields, respectively (Brun et al., 2008; Giocomo et al., 2014). In our simulation, each grid cell connected onto an average (rounded to integer) of 5, 12, and 16 IN cells in large-, intermediate-, and small-grid scale settings, respectively (Fig. 2). Each IN in turn connected onto an average (rounded to integer) of 195, 142, and 142 grid cells in large-, intermediate-, and small-grid scale settings, respectively. The average connectivity of the inhibitory feedback circuits between grid cells and interneurons in our model is thus 5*195=975, 12*142=1704, and 16*142=2272, for large-, intermediate-, and small-grid scales, respectively. Therefore, our simulation generally followed the dorsoventral connectivity gradient observed in the slice microcircuit (Grosser et al., 2021) in both the grid cell to IN direction and for the overall feedback loop, though not in the IN to grid cell direction. Of note, the number of IN cells was higher (3600) in our large grid scale simulations than in the medium and small scales (2500) to accommodate enough inhibition for larger CS rings. Simulation project settings included consistent neuron type population sizes (e.g., for conjunctive cells) for the modeling of the same animal (out of the 7 mice used in total) and module (e.g., large, intermediate, and small). Future work could further investigate directional connectivity in animal studies and how to further integrate that into increasingly more accurate computational models. Additional connectivity methods are described in the online documentation.

b. Parameters related to connectivity

The simulation of conjunctive cell to grid cell connection included a scaling of the Tsodyks and Markram (TM) model conductance constant parameter (g) to adjust for the number of simulated neurons compared to the full-scale mouse circuit. The scaling equation is $g_{scaled} = g * (N_s * P_f)$ where $N_s$

is the count of conjunctive cell presynaptic neurons in the simulation that could connect to a grid cell post-synaptic neuron, and $P_f$ is the full-scale probability of connectivity. Example $g_{scaled}$ values are listed in Table 1. The $P_f$ value used in the simulation is within two standard deviations of the mean probability reported by Hippocampome.org. A recent study has reported that direct connections may exist between CA1 and MEC LII cells (Butola et al., 2023), but Hippocampome.org does not yet have connection probability or synaptic signaling data for such connections. In our simulations, the place cell to grid cell connections represent an abstraction of both indirect (though intermediary layers) and possibly direct signaling. The corresponding g parameter was therefore not scaled or constrained.

The only connections that had synaptic weights applied to synaptic currents were from INs to grid cells. These weights were specific to each IN and ranged from 0.067 to 1.0 to fit the CS distribution. The peak weight values of 1.0 corresponded to weights not affecting signaling that is derived from g values consistent with Hippocampome.org for these neuron type pairs. These weights did not change during the simulation and can be conceptually considered to result from learning via long-term plasticity during the animal's development. Simulating this learning process was beyond the scope of this work but could be studied in future work.

V. Movement control

a. Speed Control

Network signaling levels were adjusted based on the experimentally measured speed of the real animal at each timestep (for more specific developer documentation, see Github.com/Hippocampome-Org/spatial_nav). The speed control function was optimized to perform best at speeds the animal was observed to have 98% of the time. The remaining 2% of speeds were considered outlier high speeds and the function was not optimized for them. These speed percentages are based on evaluating 10 animal experiments with a combined recording time of 12.8 hours and include the data sets used to create all results in Fig. 2. This speed control function is combined with automated direction signaling to control bump movements. The direction signaling uses a movement path function, e.g., direction angles from animal recordings, to automatically control which grid cells receive conjunctive cell input. The preferred direction (PD) of each grid cell influences the bumps to move in that same direction. Specifically, speed control of bump movement was controlled by combining conjunctive cell signals that were specific to a PD with those not specific to a preferred direction (NPD). Effective signal level combinations were identified through a parameter sweep of PD and NPD signals to test their effects on bump speeds. PD signals were fitted with a polynomial regression function and NPD signals were typically fit with sigmoidal functions, though in some cases NPD signals were simply set to a constant value to obtain desired results (Lutus, 2023; MyAssays Ltd., 2023). The supplemental data site (Github.com/Hippocampome-Org/spatial_nav) has settings files that include the specific parameters used in the speed control functions. Future work could investigate how an animal learns through development how to train its speed control mechanisms.

b. Place cell input

We employed a Gaussian function for each place cell to help control grid cell firing, and consequently bump movements, as in earlier models (Solanka et al., 2015). A balance was found through experimentation of a level of place cell firing that helped correct for drift without causing an excessive firing rate difference in the bump it affected compared to other bumps. Real animal physical space plots often have differences in firing between fields. A field with greater firing in real data plots was chosen as a field with similarly greater firing in the simulation plots by selecting which neuron to virtually record from. For each neuron, place cell input influences which field has greater firing, which makes that selection help control the occurrence of that firing.

#### c. Grid pattern rotations

In order to reproduce the rotation typically observed in real grid patterns relative to the environmental boundaries (Stensola et al., 2015), our simulations added a corresponding angle to the animal movement path signal from conjunctive cells to grid cells. This technique, however, also affects how grid fields appear at the edges of space plots. Periodic CAN simulations arrange grid cell activities on a toroidal manifold, consistent with experimental evidence (Gardner et al., 2022). Thus, activity bumps moving past one border in neural space reappear on the opposite border. Because grid field firing in physical space reflects the movement of bumps in neural space, the wrapping around of activity bumps observed when plotting sections of a continuous manifold can generate similarly wrapped spatial firing plots. Such a wrap-around effect is not present in some real recordings (Fig. 2). To avoid undesired border wrapping effect in grid fields, our simulation adjusts the conversion factor to map animal movement trajectory in physical space to bump movement in neural space so as to keep the neural activity away from the neural borders where the wrap around occurs. The documentation available through GitHub provides more details about these approaches.

### VI. Statistical Analyses

#### a. Firing rates comparison

We compared the mean firing rates of the simulated grid cells with those of real cells as reported in Fig. 6 – source data 1 (SD1) of published work (Dannenberg, 2020) and 5 new cell recordings. Mean firing rates were calculated from physical space rate maps applying multiple functions used for plotting in SD1, such as smoothing and occupancy normalization. At least 1000 seconds of recordings were utilized from each SD1 recording.

We designed the statistical analyses of firing rates to fit the amount and diversity of data available from real cells. We first selected six archetype real cells encompassing the observed range of properties (grid scale and mean firing rate) in the 29 recorded grid cells considered here. For each archetype, we created a simulation project to produce appropriate grid pattern firing. These projects were thus designed to each recreate a real cell with grid scale and firing rate archetypically representing a section of those properties observed in real cells. The 29 real cells used in this work included more cells with a small grid scale than with a large grid scale. Small grid scale cells (N=22) were divided into high (N=11) and low firing rate (N=11) cell sets, and the large grid scale cell group (N=7) was not separated by firing rate. The groups were not further subdivided beyond these groupings because of the limited number of real cells available. We selected 1 archetype project for large scale cells and 5 for small scale cells. For each archetype group, we included a simulated grid cell that recreated the firing rate of the real cell within 0.2 Hz and appeared similar in its physical space plot through visual assessment. We then used a random number generator (MATLAB’s rand() function) to select additional grid cells. If the selected cell had a grid score above the 0.2 threshold and firing rate within the range of its corresponding real group, we included it in the comparison; otherwise, another random cell was selected, and the process repeated.

This approach recreated cell firing with grid scores above the threshold for a wide range of average firing rates with different grid scales. In particular, the simulations were able to match the highest average firing rates of real cells, along with similar grid patterns and fields, in the reference data within

0.2 Hz. Those rates were 4.3 Hz, 3.7 Hz, and 2.2 Hz, for small, intermediate, and large grid scales, respectively. Only 1 out of 29 cells had an average firing rate the simulation failed to recreate within 0.2 Hz. The cell had a large-grid scale and a 0.2 Hz average firing rate. Future work can explore suitable mechanisms to obtain a good grid score at such an especially low average firing rate given that grid scale. Possible methods to achieve this aim could include the modeling of neuromodulation, more complex connectivity or additional neuron types, and long-term plasticity. Alternatively, machine learning optimization could facilitate more extensive exploration of the parameter space to find simulation settings that fit the requirement.

b. Field size and spacing comparison

In order to measure grid field size and spacing for comparing real and simulated cells, we wrote software to automatically detect grid fields (available open-source at github.com/Hippocampome-Org/gridcell_metrics). The software detects grid fields with the use of a threshold for the amount of out-of-field firing to exclude. Out-of-field firing is approximated as firing at a location that is below a specified percentage of the peak firing level. We used a threshold of 31%, which is the value that correctly detected the most cells (25 out of 27) from our previously published data (Dannenberg et al., 2020). More details of this analysis are included in a separate article (R.G. et al., 2023). The code performs field detection with the use of spatial autocorrelogram plots. Those plots are based on the firing rate map of cell firing vs. animal position, and that map includes filtering such as occupancy normalization and smoothing (with a 3.3 cm wide two-dimensional Gaussian kernel) as previously described (Dannenberg et al., 2020). The filtering was included to facilitate comparison of the results to the published experimental plots used as source data (Dannenberg et al., 2020).

The software measures at most seven fields in the autocorrelogram plot, including the central one (relative to plot borders) and six closest fields surrounding it, to find the average field size and spacing of a grid cell. Other fields are detected by the software beyond the central seven, but excluded from analyses because they are more likely to have unwanted contact with borders and a typically hexagonal grid pattern is fully represented with seven fields. Any of the seven detected fields resulting from the erroneous merging of two or more fields was excluded from size analyses, because it would result in an artifactually inflated area. Any detected field contacting a border of the plot was also excluded from size analyses, as some of its area would be missed beyond the border. Spacing analyses did not exclude fields for these other reasons because the issues were expected to have a negligible effect on spacing measurements, and including more fields improves the measurement reliability. The software uses the DBSCAN algorithm from the Statistics and Machine Learning Toolbox of MATLAB’s (mathworks.com) to automatically find clusters of plot points, each representing a detected grid field. Grid field centroids are computed as the mean x and y coordinate values of the corresponding clusters. The area of each field is calculated as the number of pixels in the cluster. The spacing between fields is calculated as the Euclidean distance between their centroids.

To measure the scale of a grid pattern, we designed a formula to account for both field size and spacing with equal weighting. Specifically, the size and field spacing for a given neuron are first each divided by the difference between the maximum and minimum values of the corresponding measures across the population of real cells, and then summed together:
$GS_n = Si_n / (MAX(Si_r)-MIN(Si_r)) + (Sp_n / (MAX(Sp_r)-MIN(Sp_r))$,
where $GS_n$ is the grid scale for a neuron with the index n, $Si_n$ is the mean linear field size for that neuron, MAX() and MIN() are the maximum and minimum values within a set of values, $Si_r$ is the set of all linear size values for real cells, $Sp_r$ is the set of all spacing values for real cells, and $Sp_n$ is the mean field spacing for neuron n. Field linear size is computed by a diameter-like formula of $Si_n$ =

$2(Si_a/\pi)^{0.5}$, where $Si_a$ is the mean area of that neuron's grid fields. The metric is described as diameter-like because it includes the formula to find the diameter of a circle from its area. The fields are not exact circles but typically close to circular, and they are deemed close enough for the formula's use. Measuring linear field size is preferred to area to allow dimensionally consistent values with field spacing. The size and spacing measurements were converted to centimeters by multiplying them by the physical size each plotted pixel represents.

c. Rhythm-related analyses

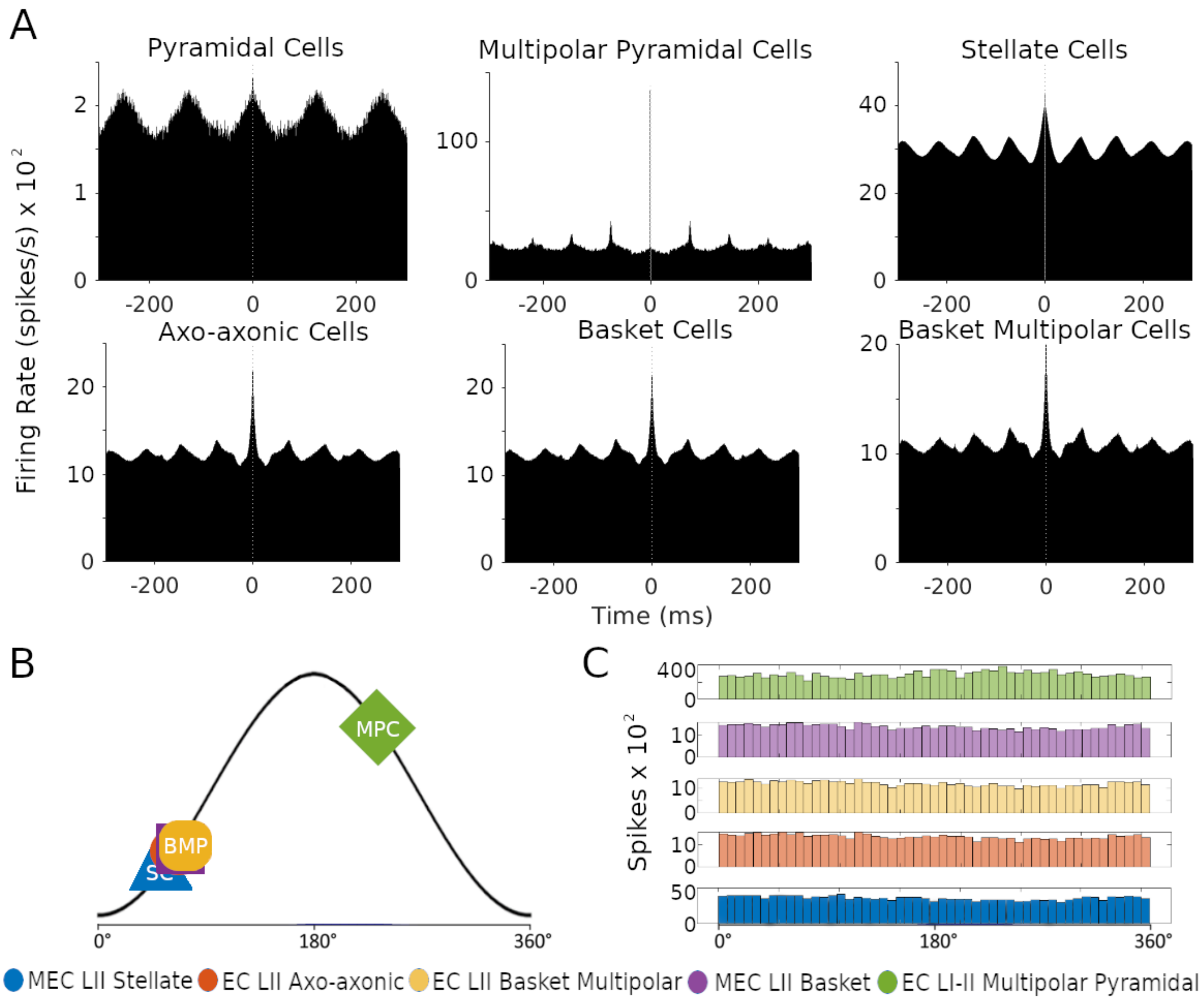


Fig. S7. Plots of rhythmic and oscillation phase activities. This activity was from the simulation in Fig. 2B. (A) The autocorrelograms of spike times for each neuron type are depicted in plots, computed from the spike times of all neurons of each neuron type. (B) Plots of the circular mean of each neuron type's phase distribution of spike times relative to the beta band spiking rhythmicity of Stellate cells. (C) Histograms display the distribution of spike-phase coupling for each neuron type.

The autocorrelations are plotted with 1-ms bin size. Normally distributed random noise with a mean of 0.01 ms and standard deviation of approximately 0.01ms was added to the autocorrelogram spike times to avoid noise in the plots due to binning artifacts. The time period used to generate all results in Fig.

S7 was the first minute of simulation time. Circular mean phases and firing rates were similar across interneuron types (Fig. S7B and S7C).

VII. Current levels

Animal experimentation has found that grid cells receive synaptic signals mediated by both fast (AMPA) and slow (NMDA) conductance (Gil et al., 2018). Specifically, the cited study reported in wild-type mice a mean of approximately 62% of total current was generated by AMPA receptors and 38% of that total generated by NMDA receptors. Another study found that mice lacking GluA1-containing AMPA receptors have impaired grid cell spatial periodicity and path integration (Allen et al., 2014), providing additional evidence toward a role of AMPA in grid cells.

The TM model parameters of Hippocampome.org represent fast conductance synaptic signaling. The same ratio of fast (AMPA) to slow (NMDA) synaptic current reported experimentally (Gil et al., 2018) was thus applied to set the corresponding conductance constant (g) values in the excitatory connections that grid cells receive. Reporting on slow current levels received by INs connected to grid cells was not found. It is assumed grid cells are influential on the synaptic current levels received in INs, given reporting of specific targeting of connections with grid cells to INs (Dhillon and Jones, 2000). It was therefore approximated that a similar ratio of fast and slow current exists on average in the grid cell synapses to INs. The simulation applied the same ratio for setting fast ($GABA_a$) and slow ($GABA_b$) conductance g constant levels for synapses from INs to grid cells because similar signal levels sent to INs are assumed to be returned from them.
